# Novel Context-Specific Genome-Scale Modelling Explores the Potential of *Chlamydomonas reinhardtii* for Synthetic Biology Applications

**DOI:** 10.1101/2022.10.07.511370

**Authors:** Herbert Yao, Sanjeev Dahal, Laurence Yang

## Abstract

Gene expression data of cell cultures is commonly measured in biological and medical studies to understand cellular decision-making in various conditions. Metabolism, affected but not solely determined by the expression, is much more difficult to measure experimentally. Thus, finding a reliable method to predict cell metabolism for given expression data will greatly benefit model-aided metabolic engineering. We have developed such a pipeline that can explore cellular fluxomics from expression data, using only a high-quality genome-scale metabolic model. This is done through two main steps: first, construct a protein-constrained metabolic model by integrating protein and enzyme information into the metabolic model. Secondly, overlay the expression data onto the modified model using a new two-step non-convex and convex optimization formulation, resulting in context-specific models with optionally calibrated rate constants. The resulting model computes proteomes and intracellular flux states that are consistent with the measured transcriptomes. Therefore, it provides detailed cellular insights that are difficult to glean individually from the omic data or metabolic models alone. As a case study, we apply the pipeline to interpret triacylglycerol (TAG) overproduction by *Chlamydomonas reinhardtii*, using time-course RNA-Seq data. The pipeline allows us to compute *C. reinhardtii* metabolism under nitrogen deprivation and metabolic shifts after an acetate boost. We also suggest a list of possible ‘bottlenecking’ proteins that need to be overexpressed to increase the TAG accumulation rate, as well as discussing other TAG-overproduction strategies.

## Introduction

Microalgae have long been a promising class of organism as a synthetic biological chassis due to their high growth rate, efficient photosystem, and simplicity in cultivation. Because of the carbon fixation capability, it is also believed that algal products have less carbon dioxide emissions and are more sustainable in large-scale production (***Khan and Fu, 2020***). As one of the best-studied microalgae, *Chlamydomonas reinhardtii* has been utilized to make a wide variety of chemicals in the lab. On the high value-added end, non-native proteins are expressed and produced in *C. reinhardtii* as pharmaceutics such as vaccines, antibiotics, and nutritional supplements (***Scaife et al., 2015***). This has been the more economically profitable and fruitful direction, and some standard workflows and toolkits have been established (***Crozet et al., 2018***). On the other hand, efforts are made to produce biofuels such as biodiesel, biohydrogen, and bio-alcohol from algae, which are chemicals closely related to the primary metabolism (***Scranton et al., 2015***). Some remarkable progress is made that increases *C. reinhardtii* lipid and starch contents by up to 2.5-fold by relatively simple modifications (***Khan and Fu, 2020***). For example, ***Rengel et al. (2018)*** showed that overexpressing the acetyl-coenzyme-A (acetyl-CoA) synthetase gene can achieve up to 2.4-fold triacylglycerol (TAG) than the control group. Some sophisticated studies have achieved an 8-fold hydrocarbon increment from the controlled *C. reinhardtii*, using gene knock-out, heterologous expression, and triparental conjugation techniques (***Yunus et al., 2018***).

Despite these findings, algal starch derivatives, lipid derivatives, and hydrogen are not yet economically feasible substitutes for fossil fuels on the market. Most of the experimental studies focus on modifying a few genes or adding a few chemical species into the medium without dramatically changing the cell from the wild type. It is a missed opportunity, as optimizing these new strains or potentially applying them in conjunctions can achieve a much higher yield with minor added costs. The optimization usually requires quantitative measurements from the phenotype, such as RNA-sequencing, proteomic data, and extracellular metabolomics, which are available from many existing studies. Developing an in-silico workflow would greatly assist researchers in systematically understanding the cellular metabolism from phenotype measurements, which is critical to both optimize current biofuel-producing strategies and suggest novel gene targets.

Genome-scale modelling (GEM) is an in-silico tool to systematically simulate cellular expression and metabolism, which is now widely used in biotechnology and infectious disease research. GEM is species-specific, usually reconstructed by researchers from an annotated genome. Genome-scale metabolic model (M-model), the most basic yet accessible GEM that focuses exclusively on predicting metabolism by mass balance, is a linear programming problem (LP) and can usually be solved within 0.1 seconds using flux balance analysis (FBA) in COBRA Toolbox (***Heirendt et al., 2019***). However, M-model cannot directly take expression data due to lacking the gene expression and macromolecule aspect. Existing algorithms for integrating M-model with data into a context-specific M-model are centred mainly around two approaches: limiting the flux of lowly expressed reactions or defining a set of core reactions (***Opdam et al., 2017***). Most of these existing methods require users to specify parameters such as expression thresholds, making them harder to use and less accessible to a wider community.

In this study, we develop a computational pipeline, OVERLAY to address these challenges. We first formulated a protein-constrained metabolic model (PC-model) starting from the published *C. reinhardtii* M-model iCre1355 and chloroplast specific M-model iGR774 (***Imam et al., 2015***; ***Røkke et al., 2020***). On top of metabolism, PC-model has protein and enzyme concentrations as variables and can be solved using the FBA algorithm with additional benefits. Moreover, expression data from other studies were overlaid onto the PC-model for novel context-specific modelling, which can predict the respective metabolic state using FBA and flux variability analysis (FVA). Our workflow is demonstrated by Fig. 1, which consists of multiple automated algorithms. This will be especially helpful for optimizing bulk material productions from *C. reinhardtii*, and the TAG accumulation case study is done using RNA-seq data from other studies to show the efficacy of this workflow.

**Figure 1.**
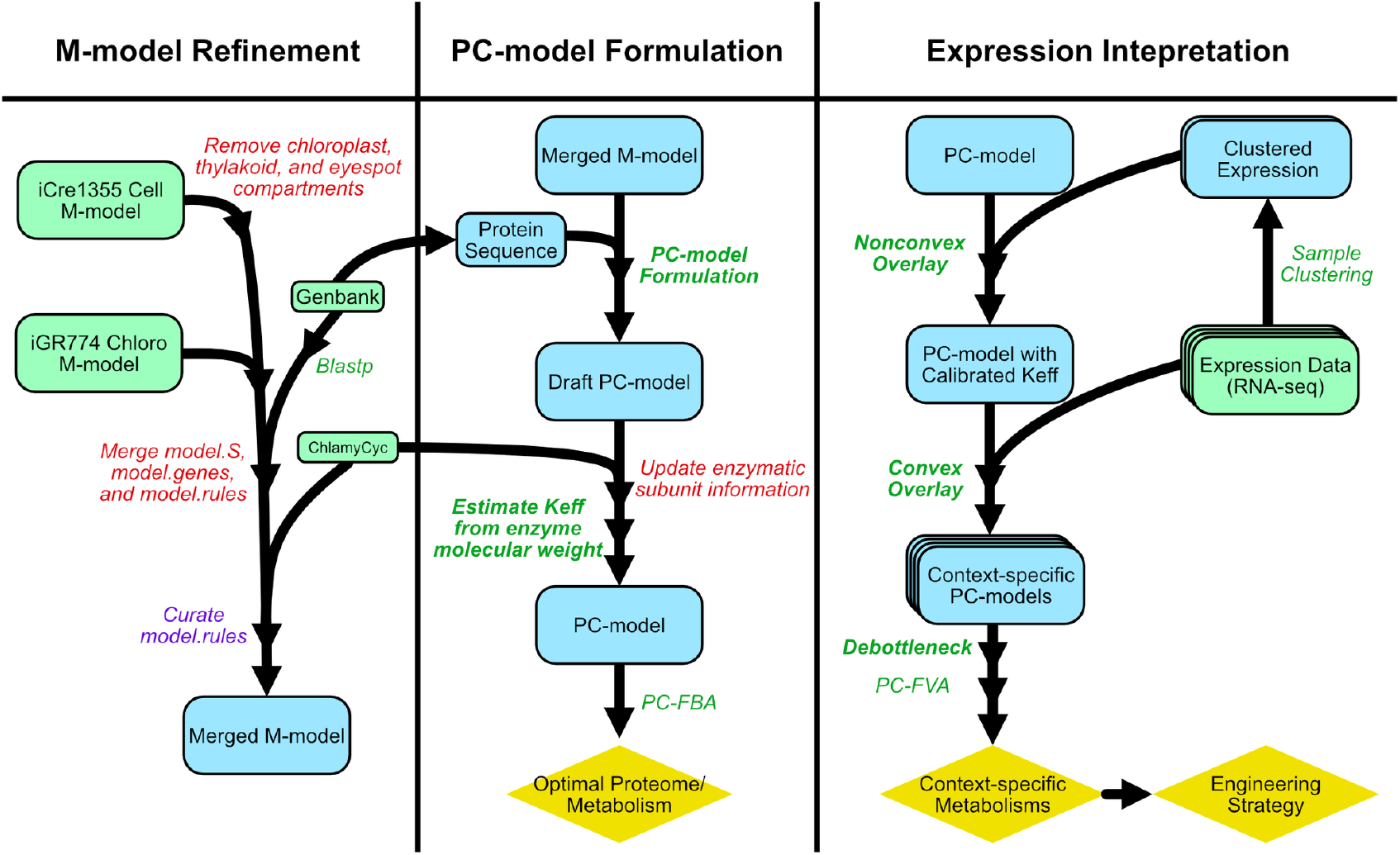
Schematic of the OVERLAY computational pipeline. Boxed texts are files and data, where green rectangle, blue rectangle, and yellow diamond denote starting materials, intermediate steps, and results, respectively. Red italic texts refer to manual procedures, and purple italic texts refer to semi-manual steps with some script aids. In contrast, automatic procedures that are done by the only script are shown in green italic texts, and methods developed in this study are in bold green font.

## Results

### Refined PC-FBA Reveals Optimal Chloroplast Metabolism, Cellular Metabolism, and Transportation

We consider *C. reinhardtii* PC-model to be a superior version of the basic M-model, as it can be used for simple analyses such as FBA with better accuracy and more utilities. Our PC-model formulation is similar to ***Yurkovich et al. (2019)***, adding protein concentrations and enzyme concentrations as variables into the model, which constrains respective reaction fluxes. The total proteome budget has defaulted to 150 mg per gram of dry cell weight (gDW).

A noticeable advantage of PC-model is that exchange reaction boundaries do not need to be set manually. This allows accurate phenotype simulation without uptake flux measurements, thus further offering a convincing comparison of flux networks between different metabolic modes. For example, by assuming sufficient lighting, photon exchange lower bound can be opened to −1000 mmol/gDW/h for any metabolic mode, and the exchange fluxes are solved by the respective optimization. Only the mixotrophic acetate uptake lower bound is manually constrained to −2 mmol/gDW/h to mimic a limited substrate availability.

We used PC-model to simulate the optimal growth strategy in autotrophic, mixotrophic and heterotrophic conditions. We compute metabolic shifts between these conditions focusing on chloroplast metabolism and transportation and interactions with the mitochondria (Fig. 2a). PC-FBA enabled us to compute the proteome allocation corresponding to the flux state in each condition (Fig. 2b-d). Photosynthesis-associated proteins accounted for 43.2 to 80.2% of proteome mass in all three conditions (Fig. 2b-d). Correspondingly, photosynthesis largely drives shifts in overall flux states (Fig. 2a).

**Figure 2.**
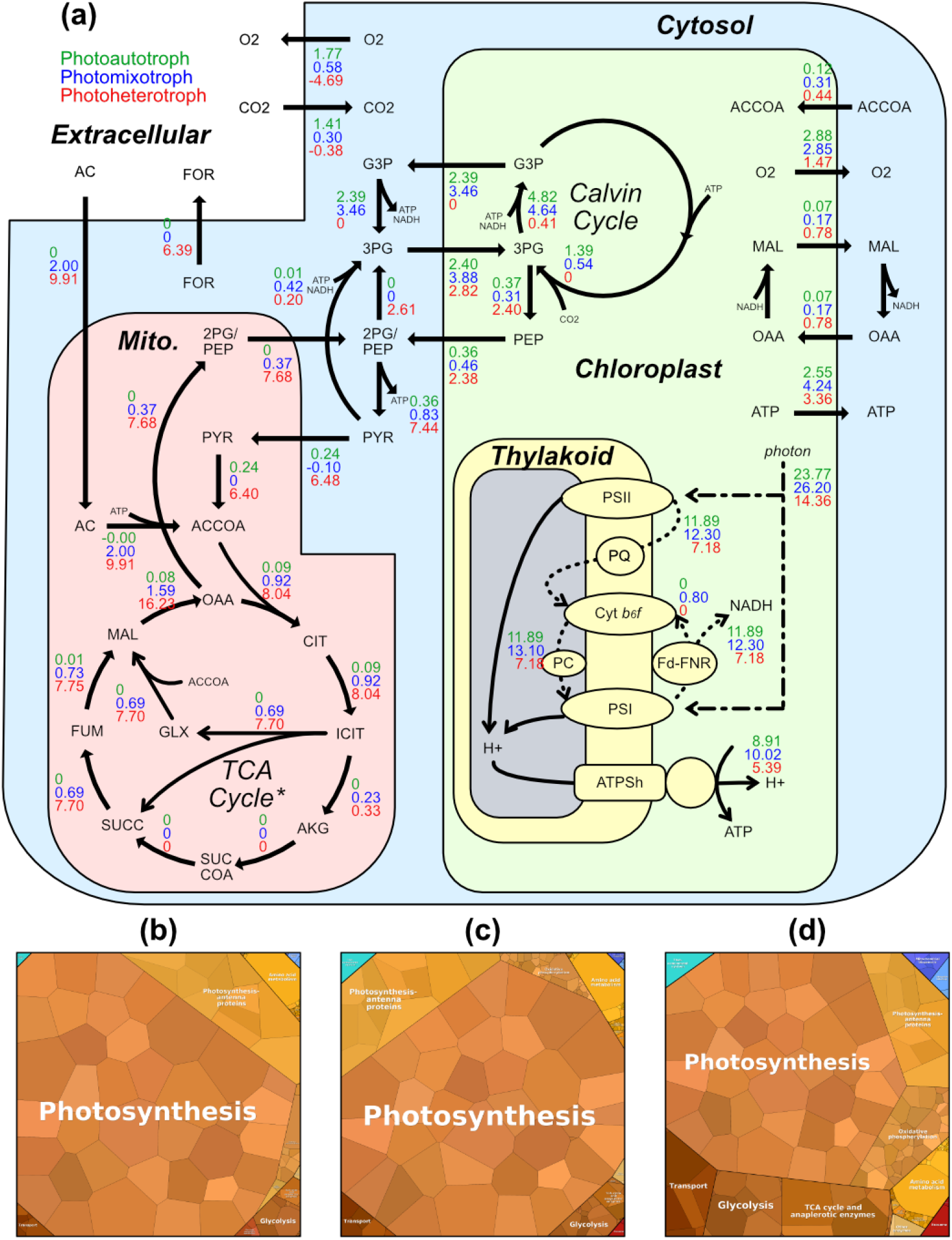
PC-FBA simulation results of autotrophic, mixotrophic, and heterotrophic *C. reinhardtii* growth mode. The optimal metabolic fluxes are shown in (a), where autotrophic, mixotrophic, and heterotrophic fluxes are denoted by green, blue, and red numbers, respectively. All fluxes are shown in mmol/gDW/h. A negative flux value means the flux is flowing in the opposite direction of the arrow. Dotted lines show the electron flux. All metabolites are shown in BiGG ID (***Norsigian et al., 2020***). Complex/enzyme abbreviations in thylakoid: PSII, photosystem II; PQ, plastoquinone/plastoquinol; Cyt b6f, cytochrome b6f complex; PC, plastocyanin; PSI, photosystem I; Fd, ferredoxin; FNR, ferredoxin NADP+ reductase; and ATPSh, CF0F1 ATP synthase. The optimal proteome of three growth modes is shown by proteome maps of (b), (c), and (d) (***Liebermeister et al., 2014***). Each small polytope is a single protein, and its area denotes the relative abundance. Likely coloured polytope is classified under the same subsystem, which is written in white text. Only modelled proteins are drawn. *Some reactions of the TCA cycle are outside of mitochondria.

Our model qualitatively reproduced major and subtle shifts determining the optimal electron flow through photosystems I and II in different conditions. *C. reinhardtii* thylakoid can choose between circular electron flow (CEF) of photosystem I only and linear electron flow (LEF) through both photosystem II and photosystem I, while LEF is the more energy-efficient option (***Arnon et al., 1958***). Consistent with this knowledge, the autotroph and heterotroph utilized LEF exclusively to maximize efficiency, as seen by zero flux from Fd to Cyt-b6f (the circular step of CEF) (Fig. 2a). Meanwhile, the mixotroph utilizes CEF, diverting 6% (0.80 mmol/gDW/h out of 13.10 mmol/gDW/h) of electron flux away from NADH/NADPH production back into the CEF. This optimal flux state suggests that CEF can result in faster growth over pure LEF under certain conditions. The result may explain how CEF and LEF are used to balance ATP/reducing power ratio for carbon fixation, as reported by ***Chaux et al.(2015)***.

Another observation is that the mixotroph has a higher photosystem activity than the other growth modes: 13.10 mmol/gDW/h electron flux from Cyt-b6f to PC for mixotroph compared to 11.89 and 7.18 mmol/gDW/h for autotroph and heterotroph, respectively (Fig. 2a). These differences are explained by PC-FBA. The autotroph needs a large portion of the proteome budget to conduct other anabolism such as carbon fixation and gluconeogenesis. For the heterotroph, it is more optimal to spend the proteome budget on consuming acetate, which serves as both an organic carbon source and an energy source. The mixotroph has limited acetate that might be enough for organic carbon but insufficient as an energy source, thus having the most potent photosystem to harvest energy.

PC-FBA also offers insightful chloroplast metabolism and transport simulations, especially regarding carbon fixation and triose-phosphate transport. As verified by other studies, 9 moles of ATP and 6 moles of NADH are required to produce 1 mole of triose-phosphate from carbon dioxide through the Calvin cycle (***Johnson and Alric, 2013***). Due to this high energy consumption, carbon fixation appears to be a suboptimal growth strategy compared to acetate uptake and is only active when acetate supply is insufficient.

Under heterotrophic growth, the chloroplast is a net consumer of organic carbon, which is transported as 3-phosphoglycerate. Noticeably, the chloroplast in all phototrophic modes uptakes 3-phosphoglycerate while excreting other triose-phosphate (Fig. 2a). Being the energy supplier of the cell, autotrophic and mixotrophic chloroplast mainly excretes the energy-compact glyceraldehyde-3-phosphate, which has been a phenomenon reported by other studies (***Johnson and Alric, 2013***). The mixotroph has the most active chloroplast, exporting more ATP and reducing power in the form of glyceraldehyde-3-phosphate and oxaloacetate/malate exchange. In all conditions, the chloroplast also consumes various amino acids while producing lipid precursors and six-carbon sugars, which are not shown in the figure in detail.

Our PC-model provides valid mitochondrion metabolism and cellular exchange simulations for all growth modes. The mixotroph and heterotroph simulations used the glyoxylate shunt in the mitochondria (Fig. 2a), as reported previously by ***Johnson and Alric*** (***2013***). This process generates excess oxaloacetate for heterotroph, which is converted to phosphoenolpyruvate and exported to the cytosol from the mitochondria. Additionally, the heterotroph excretes formate, which is only present when proteome constraints are applied (see Table S1). Thus, the proteome constraints are required to correctly predict respiro-fermentation, or overflow metabolism, as observed in multiple organisms including *C. reinhardtii* (***Gfeller and Gibbs, 1984***; ***Basan et al., 2015***). In particular, the optimal heterotroph allocates 43.2% of proteome mass to photosynthesis (Fig. 2d), compared to 80.2% in the mixotroph (Fig. 2c). The reduced photosynthesis protein budget in the heterotroph is allocated instead partially toward glycolysis and TCA cycle proteins (total 15.0%) (Fig. 2d).

Meanwhile, the autotroph shows several contrasting metabolic activities to the heterotroph. The autotrophic mitochondrion has very little activity, mostly powered by importing pyruvate from the cytosol. Instead, it generates phosphoenolpyruvate in the chloroplast, which is transported to the mitochondrion (Fig. 2a). The autotroph does share some characteristics with the mixotroph such as consuming carbon dioxide and producing oxygen. These PC-FBA simulations suggest that the mixotrophic flux network is more well-balanced and might be the ideal candidate for metabolic engineering.

### Investigating Lipid Accumulation in Nitrogen-Deprived *C. reinhardtii*

Based on insights gained from PC-FBA simulations, we further investigated mixotrophic conditions for bulk metabolite overproduction. The accumulation of TAG, a useful and value-added industrial compound, has been studied extensively in *C. reinhardtii* by inducing nitrogen deprivation. In particular, ***Goodenough et al.*** (***2014***) investigated a *sta6* (unable to form starch) strain of *C. reinhardtii* and showed enhanced TAG accumulation under nitrogen deprivation with acetate boosting 48 hours later. The study collected time-course RNA-Seq over four days of culture and discovered highly complex gene expression dynamics: 425 genes up-regulated and over 850 genes down-regulated in response to acetate (***Goodenough et al., 2014***). Here, we use our OVERLAY to decipher how these complex gene expression dynamics drive flux changes that ultimately lead to enhanced TAG production.

OVERLAY finds context-specific fluxomes and proteomes with calibrated rate constants We first directly used convex QP to overlay each of the 16 time-course samples onto the PC-model, resulting in 16 context-specific PC-models. Importantly, enzymatic rate constant (*K*_eff_) values are difficult to determine because most values are not experimentally available. We assume initially that *K*_eff_ are centred around a basal value of 65*s*^−1^, and they are proportionally scaled to the SASA (***O’brien et al., 2013***; ***Lloyd et al., 2018***) (Methods). Across all samples, the best-fitted proteome vectors are consistent with RNA-seq data, with *R*^2^ ranging between 0.950 and 0.963, with a median *R*^2^* = 0.958 (see Figure S1a for the complete plot). For example, sample 4 (time = 4 hours) has *R*^2^* = 0.954, with 56 outliers (≥ 3 times inconsistency) and 16 far outliers (≥ 10 times inconsistency) out of 1495 proteins (Fig. 3a).

**Figure 3.**
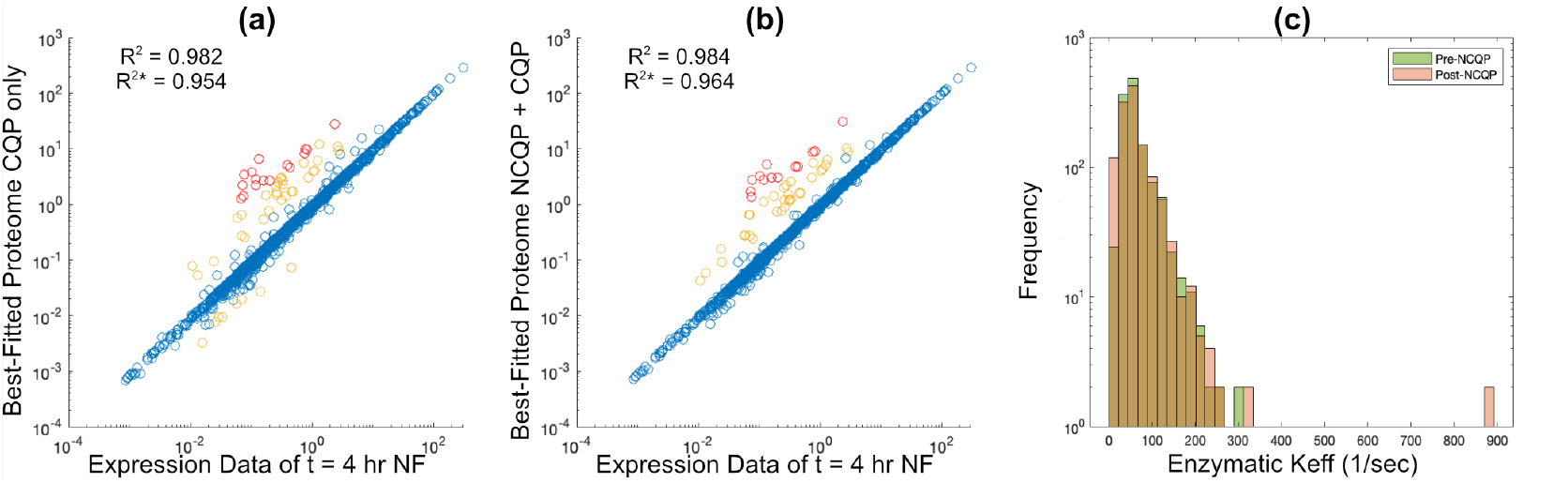
Consistency of simulated proteomes to transcriptomics, and estimated rate constants. Best-fitted proteome versus measured transcriptomes at t = 4 hours before (a) and after (b) the enzymatic rate constant adjustment by non-convex QP. *R*^2^* is computed using log-transformed data for simulated proteomes and measured transcriptomes, whereas *R*^2^ is computed without log-transforming. Outliers are denoted in yellow (≥ 3 times) and red (≥ 10 times). *R*^2^ values are computed using all points, including outliers. (c) Model *K*_eff_ values before calibration using OVERLAY (pre-NCQP) and after calibration (post-NCQP). Here, 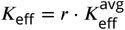.

Our OVERLAY optimally tuned the *k*_eff_, or equivalently *r*, to achieve the best fit of simulated proteomes to the RNA-Seq, subject to carefully formulated constraints (see Methods). We clustered 16 RNA-seq samples into four groups (Figure S2), which were used to estimate a single *r* vector representing all samples. This results in an improved best-fitted proteome from the original with a median *R*^2^* = 0.966 across 16 samples (Figure S1b). Fig. 3b has 48 outliers, 15 far outliers, and a higher *R*^2^* than using convex QP only. Noticeably, the fitting improvement is achieved by varying *r* only slightly from *r_ori_*. According to Fig. 3c, the distributions of *r* before and after nonconvex QP adjustment are similar, although a few enzymes are assigned much higher *r* than before. Only 214 out of 1222 enzymes have a *r* different from *r_ori_* due to extra constraints on OVERLAY (Methods, Eq. (18)), which are placed to reduce the number of total adjustments to *r*.

Additionally, OVERLAY helps to quality control the metabolic reconstruction, especially regarding its gene-reaction association. We identified a set of proteins whose abundances could not match measurements across all samples. Because we allowed for adjusted rate constants, we hypothesized that the reason for these inconsistencies is due to mis-annotations in the original model reconstruction. We manually inspected all 15 proteins that are far outliers in at least 8 out of 16 samples and compared them with their annotated functions in ChlamyCyc and ALGAEPATH (see Table S2a for the full list and details) (***Hawkins et al., 2021***; ***Zheng et al., 2014***). Indeed, we found that 5/15 proteins had incorrect gene-protein-reaction associations in the reconstruction (see Table S2a). Of the remaining 10 inconsistent proteins, we found potential isozymes for four proteins. We found a total of eight potential isozymes (Table S2b), which are promising candidates for future studies.

#### Context-Specific PC-models Providing New Metabolic Insights

The main merit of the context-specific PC-model is converting expression data to metabolic fluxes, which are insightful both independently and comparatively across samples. For example, the maximum TAG production rate is slightly reduced by ammonium-free medium and slightly promoted by the acetate boost (Fig. 3d), yet it does not translate to the ‘actual’ accumulation rate, as any point on the bar is possible for *C. reinhardtii* to operate on.

Using the final PC-FBA model, including calibrated *k*_eff_, we performed a systems-level analysis of dynamic shifts in the proteome and fluxome for mixotrophic TAG production. Given the best-fit proteome for every RNA-Seq sample, we computed the corresponding fluxome using protein-constrained flux variability analysis (PC-FVA) with TAG production rate constrained to ≥ 0%, 50%, 90%, and 99% of the maximum (Fig. 4a). For each reaction, we computed the Spearman rank correlation (*ρ*) between the max flux from PC-FVA and total abundance of all transcripts associated with the reaction. From this procedure, among all 1876 enzymatic reactions, we classify 130 reactions as expression-dependent (*ρ* ≥ 0.8), 218 reactions as expression-correlated (0.5 ≤ *ρ* < 0.8), and 1528 reactions as expression-independent (*ρ* < 0.5), while spontaneous reactions are always expression independent (Figure S3).

**Figure 4.**
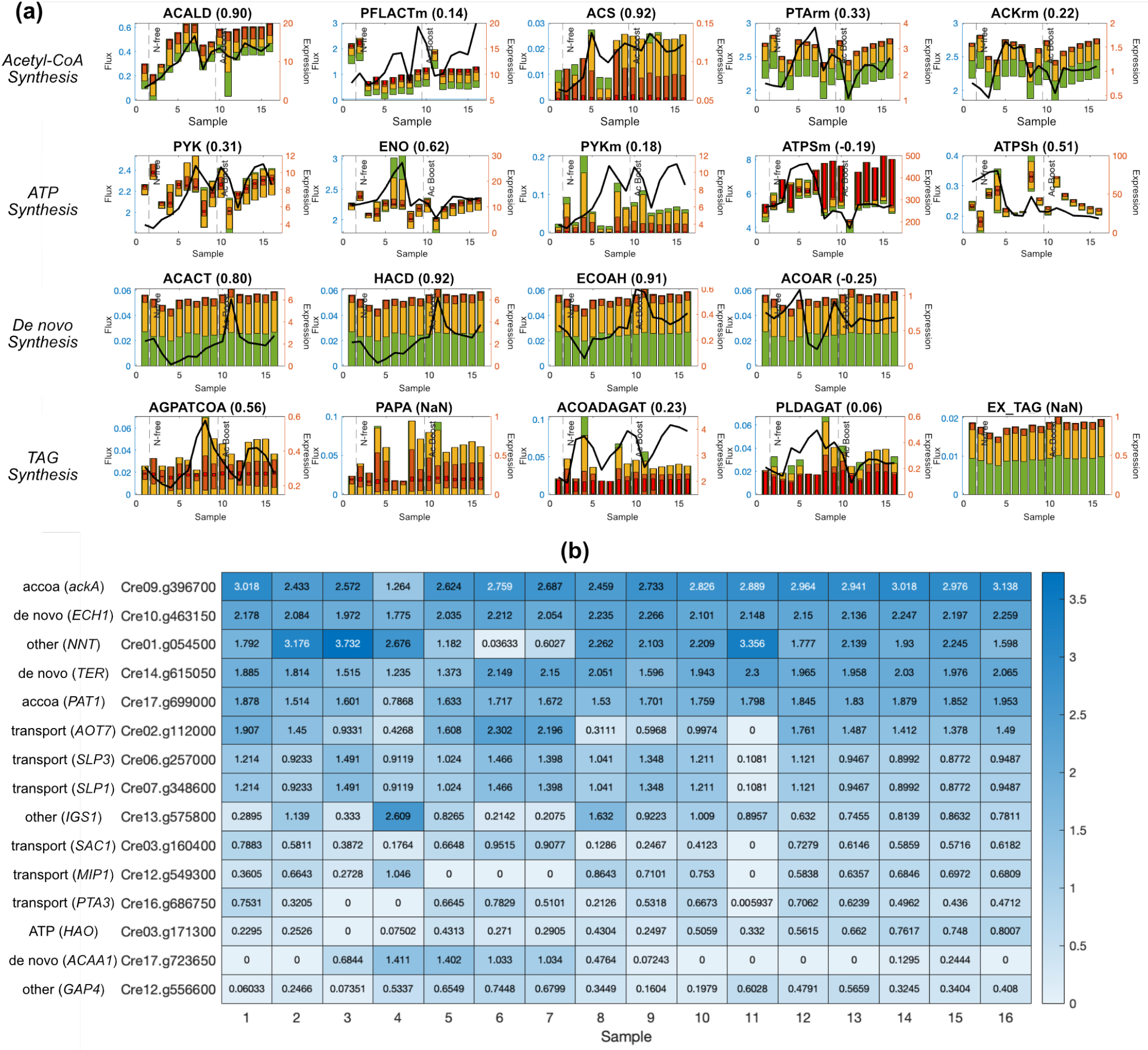
Various results of expression data interpretation. (a) is a selected collection of PC-FVA results across 16 samples regarding acetyl-CoA synthesis, ATP synthesis, De novo synthesis, and TAG synthesis pathway reactions. The bar plot shows the variability of each metabolic reaction and is coupled to the left y-axis in mmol/gDW/h. Green, yellow, orange, and red bars reflect the flux variability at 0%, 50%, 90%, and 99% of maximum TAG synthesis rate (i.e., EX_TAG flux), respectively. The black line plot shows the expression level of the reaction and is coupled to the right y-axis. In the case of isozymes presence, the black line plots the numerical sum of all isozyme levels. The calculated Spearman’s rank correlation (*ρ*) between the FVA result and expression is listed in the bracket. Vertical dashed lines divide time-series samples into pre-wash, N-free, and post-boost phases. (b) shows the amount of protein concentration (nmol/gDW) added to 15 bottleneck proteins to maximize TAG production while minimizing deviation from the measured RNA-Seq (through the protein abundance constraints). Gene symbols and functional categories of each protein are shown to the left of each gene ID (row labels).

From these PC-FVA results, especially with high optimum percentages (orange and red bars in Fig. 4a), we find that the key reactions for TAG production can be categorized into acetyl-CoA synthesis, ATP synthesis, *De novo* synthesis of free fatty acids, and TAG synthesis (Fig. 4a).

##### Acetyl-CoA Synthesis

The majority of acetyl-CoA was supplied from acetaldehyde dehydrogenase (ACALD), formate C-acetyltransferase (PFLACTm), and phosphotransacetylase (PTArm). Of these reactions, ACALD is the only expression-dependent (Spearman rank *ρ* = 0.9) reaction (Fig. 4a). ACALD is associated solely with the gene Cre17.g746997, and shows high expression correlation even for PC-FVA computations with TAG production ≥ 99% of the maximum (Fig. 4 red bars). Thus, TAG production is strongly dependent on ACALD flux, which in turn is strongly expression-dependent. Thus, Cre17.g746997 is an over-expression candidate to increase acetyl-CoA supply.

On the other hand, PFLACTm and PTArm fluxes are uncorrelated with gene expression (Spearman rank *ρ* < 0.5). PFLACTm, producing acetyl-CoA by converting pyruvate to formate, is likely dictated by the upstream pyruvate mass balance. PTArm catalyzes the highest flux of acetyl-CoA production and is dictated by both its own (Cre09.g396650 or Cre17.g699000) expression and acetate kinase (ACKrm) (Cre09.g396700 or Cre17.g709850) expression, assuming sufficient acetate supply from the medium. Over-expressing these genes will most likely boost the system-level acetyl-CoA availability. Acetyl-CoA synthetase (ACS) produces acetyl-CoA using acetate like PTArm, and it also coincides with the expression. However, its flux is nearly 100-fold lower than PTArm. Furthermore, maximum TAG flux is achieved when ACS flux is low while PTArm flux is high (Fig. 4a). This result suggests that ACS flux can be controlled through gene expression and that to maximize TAG production its expression should be repressed.

##### ATP Synthesis

ATP production is dominated by cytosolic pyruvate kinase (PYK) and mitochondrion ATP synthase (ATPSm), while the photosystem (ATPSh) activity is relatively low across all samples. This mode of ATP synthesis observed for TAG production contrasts sharply with PC-FBA simulations of optimal growth that showed large proteome allocation toward photosystems I and II (Fig. 2a). The acetate boost starts from sample 10—prior to the boost, ATPSh flux shows moderate variability and low flux relative to mitochondrial ATPS. After the acetate boost, ATPSh flux variability is even narrower, and TAG production is maximized with lowered ATPSh flux. The good correlation of ATPSh flux with gene expression (Spearman rank *ρ* = 0.51) indicates that *C. reinhardtii* is programmed to reduce photosystem-based ATP synthesis under mixotrophic growth with nitrogen limitation. Indeed, the dynamic regulation of ATP production between mitochondria and chloroplast has been studied (***Bulté et al., 1990***).

PYK produces ATP by consuming phosphoenolpyruvate. PYK flux and protein allocation differs notably between the optimal mixotrophic and heterotrophic flux networks (2PG/PEP to PYR in the cytosol in Fig. 2A). During TAG production, PYK flux is not strongly correlated with gene expression, indicating that other constraints such as mass balance determines its flux. Specifically, the primary substrate of PYK, phosphoenolpyruvate is produced by enolase (ENO). ENO is correlated with expression of Cre12.g513200 (Spearman rank *ρ* = 0.62). In turn, PYK flux is highly correlated with ENO (Spearman rank *ρ* = 0.994); therefore, both PYK and ENO flux can be controlled by regulating the Cre12.g513200 gene.

##### *De novo* free fatty acid and TAG synthesis

An intuitive way to maximize TAG production would be to overexpress its direct biosynthesis genes. This strategy has been applied for multiple algae species, including *C. reinhardtii* (***Iwai et al., 2014***). TAG is synthesized from glycerol 3-phosphate by a sequence of four reactions: 1-hexadecanoyl-sn-glycerol 3-phosphate O-acyltransferase (AGPATCOA), phosphatidate phosphatase (PAPA), acyl-CoA diacylglycerol acyltransferase (ACOADAGAT), and phospholipid diacylglycerol acyltransferase (PLDAGAT). Three acyl groups are attached in the process, using acyltransferase reactions (AGPATCOA, ACOADAGAT, PLDAGAT). TAG accumulation has been increased by overexpressing AGPATCOA in *Cyanidioschyzon merolae* (***Fukuda et al., 2018***), PLDAGAT in *C. reinhardtii* (***Iwai et al., 2014***) and Phaeodactylum tricornutum (***Zhang et al., 2021***). These strategies work by pulling carbon flux to TAG.

Our simulations are consistent with these observations, in that maximum TAG accumulation requires elevated expression of the acyltransferase proteins. Namely, the minimum required flux of AGPATCOA was above ~ 0.01 to achieve max TAG flux (Fig. 4a–*TAG synthesis*). The two diacylglycerol acyltransferase (ACOADAGAT and PLDAGAT) are also required to maximize TAG flux but because either reaction can be used, each flux has a minimum requirement of zero.

Our simulations indicate that another, possibly more critical, strategy for TAG production is to provide ample acyl-CoA by overexpressing free fatty acid synthesis cycle reactions: ACAAcetyl-CoA C-acyltransferase (ACACT), 3-hydroxyacyl-CoA dehydrogenase (HACD), and enoyl-CoA hydratase (ECOAH), and trans-2-enoyl-CoA reductase (ACOAR). These reactions showed time-course flux patterns that were nearly identical to the TAG dilution reaction (EX_TAG) (Fig. 4a). Three of these reactions (ACACT, HACD, and ECOAH) are highly correlated with gene expression (Spearman rank *ρ* = 0.8, 0.92, and 0.91). Therefore, over-expression of these genes would directly increase flux. Indeed, a sharp over-expression of these genes coincides well with the acetate boost (Fig. 4a, *De novo synthesis*). Finally, the ACOAR reaction is uncorrelated with gene expression (Spearman rank *ρ* = −0.25); however, due to mass balance constraints, its flux is entirely determined by the flux of the preceding three reactions in the cycle.

#### Engineering Strategies for Enhancing TAG Productivity

Next, we developed a tool to find optimal, system-level debottlenecking strategies for metabolic engineering. This tool is formulated as a linear program (Methods): TAG production is maximized subject to all context-specific PC model constraints, while also allowing for a user-defined total protein over-expression “budget” (*E*). The optimal solution to this problem provides the set of highest priority targets for protein overexpression.

Using this tool, we identified protein overexpression strategies that were consistent with the transcriptome changes observed in the acetate-boosted experiment. For example, all three free fatty acid synthesis genes (*ECH1, TER, ACAA1*) that were highly correlated with reaction flux (ECOAH, HACD, ACACT) were identified as overexpression targets (Fig. 4b).

Additionally, two of the acetyl-CoA synthesis genes (*ackA and PAT1*), corresponding to ACKrm and PTArm reactions were identified as overexpression targets. Interestingly, the algorithm did not identify the gene for ACALD as an overexpression target, despite it being a key step in Acetyl-CoA synthesis. This result is consistent with the high expression levels of ACALD-associated transcripts (Fig. 4a, ACALD); therefore, no further debottlenecking is required. In fact, this result suggests that the expression levels for PATrm and ACKrm-associated genes may need to be increased further to achieve TAG production higher than that observed.

Finally, we identified additional overexpression candidates that may be candidates for future engineering. These proteins include six transporters, including those for sulphate (*SLP1, SLP3*), phosphate (*PTA3*), and amino acids (*AOT7*) (Fig. 4b). Other targets include proteins associated with amino acid biosynthesis (*IGS1*), glycolysis (*GAP4*), redox balance (*NNT*), and glyoxylate metabolism (*HAO*) (Fig. 4b).

Another possible TAG productivity-enhancing approach is to supplement the medium with additional carbon sources. The measured transcriptomes indicated the expression of transporters for alternative carbon substrate uptake including acetamide, D-ribose, and D-lactate. By adding these carbon sources to the *in silico* growth media, we confirmed that the observed gene expression levels support uptake of these carbon sources, albeit at slow rates. Our simulations indicate that supplementing these carbon sources would not boost TAG production under these conditions because the added carbon would not alleviate the main bottleneck, free fatty acid biosynthesis.

## Discussion

In this work, we developed a computational pipeline for building a context-specific, protein-constrained genome-scale model (PC-model), starting from metabolic reconstruction and transcriptomics data. We showcase the utility of PC-model for deciphering how complex gene expression dynamics drive system-level fluxome shifts in *C. reinhardtii* using published time-course transcriptomics data. Using PC-FBA, we recapitulate metabolic hallmarks of autotrophic, heterotrophic, and mixotrophic growth. Importantly, the protein constraints are required to accurately simulate respiro-fermentation (overflow) metabolism. We then use time-course RNA-Seq data (***Goodenough et al., 2014***) to investigate the over-production of triacylglycerol (TAG) in response to acetate supplementation under nitrogen limitation. Our pipeline generated context-specific models for each experimental time-point (over four days of culture), with very high consistency between 1495 modelled proteins and measured transcriptomes (*R*^2^ between 0.95 to 0.963, median *R*^2^ = 0.958). We then determined which metabolic fluxes were controlled by gene expression. By comparing simulated fluxes and measured transcriptomes across the 16 time-course RNA-Seq samples, we could categorize all gene-associated reactions into 130 expression-dependent (Spearman rank *ρ* ≥ 0.8), 218 expression-related (Spearman rank 0.5 ≤ *ρ* < 0.8, and 1528 expression-independent (Spearman rank *ρ* < 0.5) reactions.

To enable researchers to systematically identify optimal overexpression targets, we developed a novel optimization-based tool. Using the tool, we identified key gene expression bottlenecks for TAG overproduction. The tool recapitulated known bottlenecks (e.g., the acyltransferase steps in TAG biosynthesis). Furthermore, we identified several novel overexpression targets to further improve TAG overproduction including genes encoding sulphate, phosphate, and amino acid transporters; glyoxylate metabolism, and redox balancing.

### *C. reinhardtii* as a Synthetic Biology Chassis

*C. reinhardtii* has been studied for decades as a cell factory, producing both bulk metabolites and, more recently, expressing heterologous genes for non-native value-added products. Although many established methods are available for bulk metabolite production, in some cases they are still far from optimal productivity based on our simulation results. Indeed, the recent adaptive laboratory evolution of *C. reinhardtii* has increased both growth rate (by up to 300%) and product yields (DHA production by 90%) (***LaPanse et al., 2021***). Using genome-scale modelling, especially with the context-specific PC-model pipeline, *C. reinhardtii* may become an economically-efficient cell factory for bulk chemicals after multiple iterations of optimization. On the other hand, the optimization for non-native products is more complicated, yet it can be highly impactful for the (bio)chemicals industry due to the potential for sustainable production of high value products, especially when non-biological synthesis routes are unavailable.

### Evaluation and Outlook of Context-Specific Proteome Constrained Models

Our model formulation and pipeline provides several advantages over existing methods. First, our PC-model computes optimal fluxomes in response to changes in the allocation of the proteome. This proteome, in turn, is highly consistent with measured transcriptomes through a sequence of convex and non-convex quadratic optimization problems. Second, to perform context-specific simulations, we do not require choosing an arbitrary gene expression threshold for turning on/off reactions based on transcript abundance—this has been a challenge in existing methods (***Opdam et al., 2017***). Third, our pipeline enables the identification of optimal overexpression targets. This method requires only one parameter to be adjusted: *E* (total protein overexpression budget), which can be determined using a simple procedure. Finally, our method enables using transcriptomics to quality-control genome-scale reconstructions and their annotations, which are found through persistent discrepancies between optimal protein and measured transcript abundances.

In general, the context-specific PC-model is a simple yet effective tool for understanding and manipulating cellular metabolism through gene expression, making it potentially valuable for many metabolic engineering applications. The prediction results can be used for practical decision-making in various research fields such as biotechnology, infectious disease, and cancer. Overall, we believe the context-specific PC-model is a versatile tool that will benefit the system biology and metabolic engineering community.

## Materials and Methods

### Merging iCre1355 and iGR774 Metabolic Model and Curation

Instead of using *C. reinhardtii* M-model iCre1355 only, we decided to plug in a newer chloroplast M-model, which contains more up-to-date understandings of the chloroplast metabolism. We merged M-model iCre1355 (cellular model) and iGR774 (chloroplast model) by first deleting all chloroplast metabolites and reactions in the cellular model. The chloroplast model was slightly modified (see Table S3), and its transported reactions were matched to the dead-end transportations in the cellular model (see Table S4-6). The newly merged M-model has 1354 genes, 2641 reactions, and 2240 metabolites. We curated the gene-reaction rules (stored in the model as `model.rules` field) using complex data from ChlamyCyc 8.0 (***Hawkins et al., 2021***). Only enzyme complexes with multiple subunits were curated, but not any protein-monomer enzymes (Table S7). The starch metabolism pathway and appropriate rules were also added to the merged model (Table S3). We also opened the lower bound of reaction GAPDHi and GAPDH_nadp to allow reverse reactions (Table S3). The script written to merge and modify M-models is `MergedModel.m`, which calls functions in COBRA Toolbox on MATLAB to load and manipulate M-models (***Heirendt et al., 2019***; ***MATLAB, 2021b***). We used the Kyoto Encyclopedia of Genes and Genomes (KEGG) and BiGG Models as general references in M-model modifications (***Kanehisa et al., 2021***; ***Norsigian et al., 2020***).

### Protein Constraints Implementation

Our PC-model formulation is shown below.

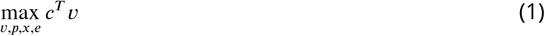

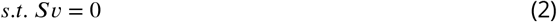

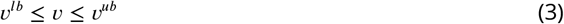

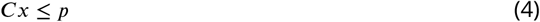

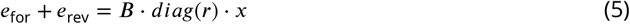

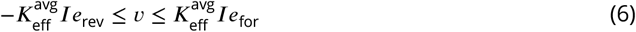

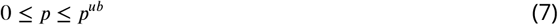

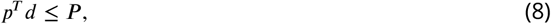

where 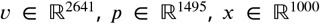, and 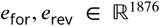 denote metabolic flux, proteome concentration, complex concentration, and enzyme concentration, respectively.

By adopting this formulation, we assumed the following:

1. The total amount of metabolic proteome may not exceed a weight fraction of the dry weight, which is further referred to as the ‘proteome budget’ and denoted by a scalar (***P***) in mg/gDW.
2. Each annotated gene in the M-model is transcribed and translated to a unique protein whose molecular weight can be estimated by its protein sequence.
3. Rate constant of a certain enzyme is fixed regardless of reactions. This will greatly reduce the complexity of the problem, especially the non-convex quadratic programming problem (non-convex QP) in the later section.
4. Enzyme concentration upper bounds but not forces the respective reaction flux. Enzymes are currently not compartmentalized.

The protein constraints were implemented in the M-model by adding four sets of variables and four sets of constraints. Variables are defined as follows:

1. Protein dilution: protein concentrations in nmol/gDW. Proteins are uniquely defined for each gene in the M-model.
2. Complex formation: complex concentrations in nmol/gDW. We define ‘complex’ as a unique protein combination that can sufficiently catalyze any single reaction. The list of complexes is obtained by parsing rules in M-model (`parseGeneRule.m`).
3. Enzyme formation: enzyme concentrations in nmol/gDW. We define ‘enzyme’ as a collection of indifferent complexes that can catalyze a certain reaction. A pair of forwarding and reverse enzymes are added for each enzymatic reaction, and no enzyme is added for spontaneous reactions.
4. Enzyme dilution: One dilution reaction for each forward or reverse enzyme.

Extra constraints were added to the model as follows:

1. Each complex may not exceed the abundance of available protein subunit, according to *C* (Eq. (4)). *C* is a matrix containing complex subunit information. Excess proteins are allowed.
2. The sum of forward and reverse enzyme equals the total complex, according to *r* and *B* (5). *B* is Boolean matrix mapping complexes and enzymes and further enzymatic reactions. Vector *r* denotes the ratio between the rate constant of each complex and the average enzymatic rate constant, or 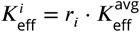. We first estimated *r* as below (`estimateKeffFromMW.m`):

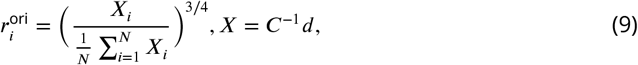

assuming larger complex generally has a larger enzymatic rate constant.
3. Enzymatic reaction fluxes are restrained by respective forward and reverse enzyme levels through the average rate constant of 65*s*^−1^.
4. Protein concentrations are collectively constrained by the proteome budget ***P*** of 150mg/gDW, according to protein molecular weight vector *d* in mg/nmol.

We collected a complete protein sequence FASTA file using NCBI genome assembly *Chlamydomonas reinhardtii* v5.5 (`GCF_000002595.2.gbff`), *Chlamydomonas reinhardtii* chloroplast reference genome (NC_005353.1), and *Chlamydomonas reinhardtii* mitochondrial reference genome (NC_001638.1) (***Sayers et al., 2021***). This FASTA was constructed by extracting all locus tags and respective protein sequences into a plain text file (`fastaParsing.m`). It was used to calculate the molecular mass of modelled proteins (`calcProteinMM.m`). The PC-model construction processes above are also automated in a MATLAB file as `pcModel.m`. The solving time of PC-FBA is around 0.3 seconds on our device, which is 6 times more than its respective FBA.

### Obtaining and Processing Expression Data for Case Study

Raw reads of RNA-seq data (E-GEOD-56505) for the TAG case study were downloaded as FASTQ files (***Goodenough et al., 2014***). We downloaded the NCBI genome assembly `GCF_000002595.2.gbff` and parsed the GenBank file into a FASTA reference transcript (***Sayers et al., 2021***). Reads were aligned using Bowtie2 with default settings and quantified using Samtools and Salmon (***Langmead and Salzberg, 2012***; ***Danecek et al., 2021***; ***Patro et al., 2017***).

### Overlaying processed RNA-Seq Data onto PC-model using Convex QP

We proposed a methodology to interpret the underlying cellular metabolism for a given RNA-seq data using the PC-model (`overlayMultiomicsData.m`). Assuming the proteome vector is similar to the mRNA vector, we formulated a quadratic objective function, subject to constraints (2) – (8):

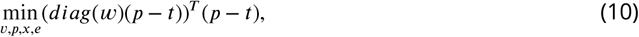

where *t* denotes the transcript abundance vector, and *ω* is a weighting vector for each transcript. This finds the proteome vector closest to the measured transcriptome while maintaining underlying metabolic feasibility. We defined

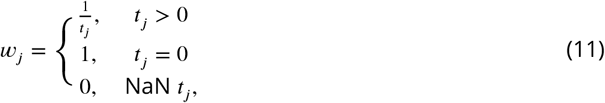

which increases the weighting of lowly transcribed and un-transcribed genes. This is essential to keep the unexpressed proteins absent from the context-specific model, although other weighting functions might be feasible too. The expression measurement was unavailable for some modelled proteins (NaN *t_j_*), such as proteins translated from the chloroplast and mitochondria genome, in which case the weighting was assigned to zero. *t* was scaled to satisfy

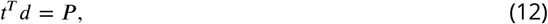

which put *t* and *ρ* into the same magnitude. This guaranteed the possibility that objective function (10) might reach zero value from solving.

This is a convex QP problem and can be solved using commercial LP solvers such as Gurobi Optimizer or IBM ILOG CPLEX (***Gurobi Optimization, 2022***; ***Cplex, 2017***). The optimization took around five seconds for both solvers. Defining the best-fitting protein vector is solved to be *p*’, and we replaced constraint (7) with constraint (13) for the base PC-model to make it sample-specific.

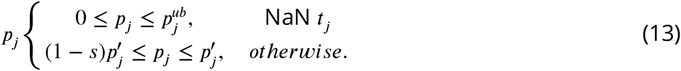

We added a slack term (*s* = 0.02) into constraint (13) for two practical reasons: leaving the proteome budget for unmeasured transcripts in the unbiased analysis and making the de-bottlenecking algorithm easier to implement. We found that eliminating the slack also indirectly restricts unmeasured proteins due to proteome budget depletion, which would impact the downstream analysis. In our practice, adjusting *s* within a reasonable range would not significantly affect the result.

### Estimating System Level Enzymatic Rate Constants using Non-Convex QP

Although it is impractical to experimentally measure all enzymatic rate constants, they can be systematically estimated using this PC-model formulation. Given the *Q* set of samples of the same strain under different conditions, we rewrote objective function (10) into objective (13). The intuition was to find the single vector *r* that allows the best-fitting result for all RNA-seq samples:

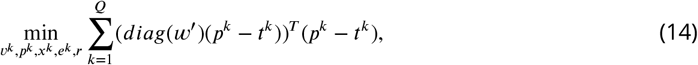

subject to constraints (2) – (8) for each of the *Q* samples. For example, constraint (2) effectively becomes

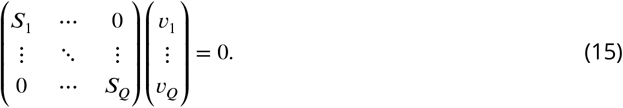

We also simplified the weighting by decorrelating *ω*’ with relative abundance, greatly speeding up the computation:

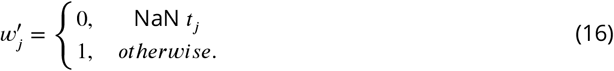

Vector *r* was made a variable with the following constraints:

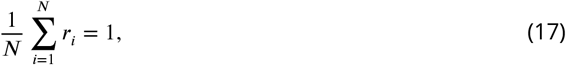

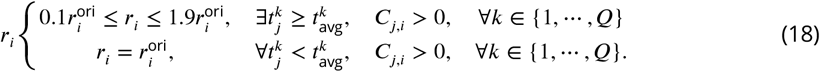

The constraints above formulate a non-convex QP that is *Q* times the size of the convex QP. By constraint (18), we preserved the original *r_i_* with low subunit abundance, which prevented the solver from prematurely modifying the rate constant. This is inferring that a more comprehensive rate constant estimation can be achieved by including data from various metabolic modes while stacking up data from similar metabolic modes will benefit little; on the other hand, adding each set of data exponentially increases the computational cost. Thus, we first performed hierarchical clustering by MATLAB Statistics and Machine Learning Toolbox to categorize 16 RNA-seq samples into four groups (Figure S2), which were then used group averages as ‘samples’ to estimate *r* using non-convex QP (***MATLAB, 2021a***). Procedures above can be done by `overlayMultiomicsData.m` with ‘keffEstimate’ option set to true.

The QP was solved by Gurobi Optimizer version 9.1.2, a state-of-the-art LP solver that supports non-convex bilinear optimizations (***Gurobi Optimization, 2022***). The optimization took around 2000 seconds on a laptop with an Apple M1 chip and 16 GB of memory, although the computation time can vary widely for the same problem size with different data samples.

### De-bottlenecking and Unbiased Network Analysis

Acknowledging errors and uncertainties in the data and workflow, we applied a de-bottlenecking optimization onto data-specific PC-models to mitigate the effect of a few bottlenecking proteins without shifting the landscape (`proteinDebottleneck.m`). This was done by adding a variable term to the constraint (13), which becomes

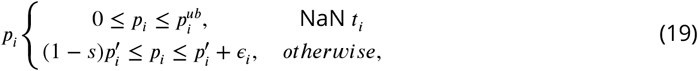

where *ϵ* is a variable vector with an assigned over-expression budget *E*:

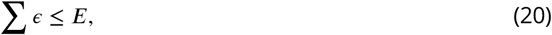

while other constraints are the same as the data-specific PC-model, optimized to objective function (1) using the FBA algorithm, referred to as protein-constrained flux balance analysis (PC-FBA). We conducted this LP by varying error budget values and eventually chose *E* = 20, where the curve of optimal objective values versus *E* reached a constant slope (see Table S8). This means no single protein was critically bottlenecking the objective function, and therefore it was a suitable state for the downstream analysis. The optimization took roughly 2.0 seconds using Gurobi Optimizer.

To further understand the metabolic capabilities under each expression data, we used PC-FVA for an unbiased analysis. For each data-specific model, FVA of all metabolic reactions *υ* was done to find *υ*_*min*_ and *υ*_*max*_ at the optimal percentages of 0%, 50%, 90%, and 99%.

## Acknowledgments

The authors thank Professor X. Li at Chemical Engineering Department, Queen’s University for his opinions and discussions on non-convex optimization; the lead author also thanks Z. Wang at Queen’s University for her general discussion.

## References

Arnon DI, Whatley FR, Allen MB. Assimilatory Power in Photosynthesis. Science. 1958 May; 127(3305):1026–1034. https://www.science.org/doi/abs/10.1126/science.127.3305.1026, doi: 10.1126/science.127.3305.1026, publisher: American Association for the Advancement of Science.

Basan M, Hui S, Okano H, Zhang Z, Shen Y, Williamson JR, Hwa T. Overflow metabolism in Escherichia coli results from efficient proteome allocation. Nature. 2015 Dec; 528(7580):99–104. https://www.nature.com/articles/nature15765, doi: 10.1038/nature15765, number: 7580 Publisher: Nature Publishing Group.

Bulté L, Gans P, Rebéillé F, Wollman FA. ATP control on state transitions in vivo in Chlamydomonas reinhardtii. Biochimica et Biophysica Acta (BBA) - Bioenergetics. 1990 Oct; 1020(1):72–80. https://www.sciencedirect.com/science/article/pii/000527289090095L, doi: 10.1016/0005-2728(90)90095-L.

Chaux F, Peltier G, Johnson X. A security network in PSI photoprotection: regulation of photosynthetic control, NPQ and O2 photoreduction by cyclic electron flow. Frontiers in Plant Science. 2015; 6. https://www.frontiersin.org/articles/10.3389/fpls.2015.00875.

Cplex II, V12. 9: User’s Manual for CPLEX. International Business Machines Corporation; 2017.

Crozet P, Navarro FJ, Willmund F, Mehrshahi P, Bakowski K, Lauersen KJ, Pérez-Pérez ME, Auroy P, Gorchs Rovira A, Sauret-Gueto, S Niemeyer J, Spaniol B, Theis J, Trösch R, Westrich LD, Vavitsas K, Baier T, Hübner W, de Carpentier F, Cassarini M, et al. Birth of a Photosynthetic Chassis: A MoClo Toolkit Enabling Synthetic Biology in the Microalga Chlamydomonas reinhardtii. ACS Synthetic Biology. 2018 Sep; 7(9):2074–2086. https://doi.org/10.1021/acssynbio.8b00251, doi: 10.1021/acssynbio.8b00251, publisher: American Chemical Society.

Danecek P, Bonfield JK, Liddle J, Marshall J, Ohan V, Pollard MO, Whitwham A, Keane T, McCarthy SA, Davies RM, Li H. Twelve years of SAMtools and BCFtools. GigaScience. 2021 Feb; 10(2):giab008. https://www.ncbi.nlm.nih.gov/pmc/articles/PMC7931819/, doi: 10.1093/gigascience/giab008.

Fukuda S, Hirasawa E, Takemura T, Takahashi S, Chokshi K, Pancha I, Tanaka K, Imamura S. Accelerated triacylglycerol production without growth inhibition by overexpression of a glycerol-3-phosphate acyltransferase in the unicellular red alga Cyanidioschyzon merolae. Scientific reports. 2018; 8(1):1–12.

Gfeller RP, Gibbs M. Fermentative Metabolism of Chlamydomonas reinhardtii. Plant Physiology. 1984 May; 75(1):212–218. https://www.ncbi.nlm.nih.gov/pmc/articles/PMC1066864/.

Goodenough U, Blaby I, Casero D, Gallaher SD, Goodson C, Johnson S, Lee JH, Merchant SS, Pellegrini M, Roth R, Rusch J, Singh M, Umen JG, Weiss TL, Wulan T. The Path to Triacylglyceride Obesity in the sta6 Strain of Chlamydomonas reinhardtii. Eukaryotic Cell. 2014 May; 13(5):591–613. https://journals-asm-org.proxy.queensu.ca/doi/full/10.1128/EC.00013-14, doi: 10.1128/EC.00013-14, publisher: American Society for Microbiology.

Gurobi Optimization L, Gurobi Optimizer Reference Manual; 2022. https://www.gurobi.com.

Hawkins C, Ginzburg D, Zhao K, Dwyer W, Xue B, Xu A, Rice S, Cole B, Paley S, Karp P, Rhee SY. Plant Metabolic Network 15: A resource of genome-wide metabolism databases for 126 plants and algae. Journal of Integrative Plant Biology. 2021; 63(11):1888–1905. https://onlinelibrary.wiley.com/doi/abs/10.1111/jipb.13163, doi: 10.1111/jipb.13163, _eprint: https://onlinelibrary.wiley.com/doi/pdf/10.1111/jipb.13163.

Heirendt L, Arreckx S, Pfau T, Mendoza SN, Richelle A, Heinken A, Haraldsdóttir HS, Wachowiak J, Keating SM, Vlasov V, Magnusdóttir S, Ng CY, Preciat G, Žagare A, Chan SHJ, Aurich MK, Clancy CM, Modamio J, Sauls JT, Noronha A, et al. Creation and analysis of biochemical constraint-based models using the COBRA Toolbox v.3.0. Nature Protocols. 2019 Mar; 14(3):639–702. https://www.nature.com/articles/s41596-018-0098-2, doi: 10.1038/s41596-018-0098-2, number: 3 Publisher: Nature Publishing Group.

Imam S, Schäuble S, Valenzuela J, López García de Lomana A, Carter W, Price ND, Baliga NS. A refined genome-scale reconstruction of Chlamydomonas metabolism provides a platform for systems-level analyses. The Plant Journal. 2015; 84(6):1239–1256. https://onlinelibrary.wiley.com/doi/abs/10.1111/tpj.13059, doi: 10.1111/tpj.13059, _eprint: https://onlinelibrary.wiley.com/doi/pdf/10.1111/tpj.13059.

Iwai M, Ikeda K, Shimojima M, Ohta H. Enhancement of extraplastidic oil synthesis in C hlamydomonas rein-hardtii using a type-2 diacylglycerol acyltransferase with a phosphorus starvation–inducible promoter. Plant biotechnology journal. 2014; 12(6):808–819.

Johnson X, Alric J. Central Carbon Metabolism and Electron Transport in Chlamydomonas reinhardtii: Metabolic Constraints for Carbon Partitioning between Oil and Starch. Eukaryotic Cell. 2013 Jun; 12(6):776–793. https://journals-asm-org.proxy.queensu.ca/doi/full/10.1128/EC.00318-12, doi: 10.1128/EC.00318-12, publisher: American Society for Microbiology.

Kanehisa M, Furumichi M, Sato Y, Ishiguro-Watanabe, M Tanabe M. KEGG: integrating viruses and cellular organisms. Nucleic Acids Research. 2021 Jan; 49(D1):D545–D551. https://doi.org/10.1093/nar/gkaa970, doi: 10.1093/nar/gkaa970.

Khan S, Fu P. Biotechnological perspectives on algae: a viable option for next generation biofuels. Current Opinion in Biotechnology. 2020 Apr; 62:146–152. https://www.sciencedirect.com/science/article/pii/S0958166919300953, doi: 10.1016/j.copbio.2019.09.020.

Langmead B, Salzberg SL. Fast gapped-read alignment with Bowtie 2. Nature Methods. 2012 Apr; 9(4):357–359. https://www.nature.com/articles/nmeth.1923, doi: 10.1038/nmeth.1923, number: 4 Publisher: Nature Publishing Group.

LaPanse AJ, Krishnan A, Posewitz MC. Adaptive Laboratory Evolution for algal strain improvement: methodologies and applications. Algal Research. 2021 Mar; 53:102122. https://www.sciencedirect.com/science/article/pii/S2211926420309905, doi: 10.1016/j.algal.2020.102122.

Liebermeister W, Noor E, Flamholz A, Davidi D, Bernhardt J, Milo R. Visual account of protein investment in cellular functions. Proceedings of the National Academy of Sciences. 2014 Jun; 111(23):8488–8493. https://www.pnas.org/doi/full/10.1073/pnas.1314810111, doi: 10.1073/pnas.1314810111, publisher: Proceedings of the National Academy of Sciences.

Lloyd CJ, Ebrahim A, Yang L, King ZA, Catoiu E, O’Brien EJ, Liu JK, Palsson BO. COBRAme: A computational framework for genome-scale models of metabolism and gene expression. PLoS Computational Biology. 2018; 14(7):e1006302.

MATLAB, Statistics and Machine Learning Toolbox. The MathWorks Inc.; 2021. https://www.mathworks.com/help/stats/.

MATLAB. version 9.11.0.1769968 (R2021b). Natick, Massachusetts, United State: The MathWorks Inc.; 2021.

Norsigian CJ, Pusarla N, McConn JL, Yurkovich JT, Dräger A, Palsson BO, King Z. BiGG Models 2020: multi-strain genome-scale models and expansion across the phylogenetic tree. Nucleic Acids Research. 2020 Jan; 48(D1):D402–D406. https://doi.org/10.1093/nar/gkz1054, doi: 10.1093/nar/gkz1054.

O’brien EJ, Lerman JA, Chang RL, Hyduke DR, Palsson BØ. Genome-scale models of metabolism and gene expression extend and refine growth phenotype prediction. Molecular Systems Biology. 2013; 9(1):693.

Opdam S, Richelle A, Kellman B, Li S, Zielinski DC, Lewis NE. A Systematic Evaluation of Methods for Tailoring Genome-Scale Metabolic Models. Cell Systems. 2017 Mar; 4(3):318–329.e6. https://www.sciencedirect.com/science/article/pii/S2405471217300108, doi: 10.1016/j.cels.2017.01.010.

Patro R, Duggal G, Love MI, Irizarry RA, Kingsford C. Salmon: fast and bias-aware quantification of transcript expression using dual-phase inference. Nature methods. 2017 Apr; 14(4):417–419. https://www.ncbi.nlm.nih.gov/pmc/articles/PMC5600148/, doi: 10.1038/nmeth.4197.

Rengel R, Smith RT, Haslam RP, Sayanova O, Vila M, León R. Overexpression of acetyl-CoA synthetase (ACS) enhances the biosynthesis of neutral lipids and starch in the green microalga Chlamydomonas reinhardtii. Algal Research. 2018 Apr; 31:183–193. https://www.sciencedirect.com/science/article/pii/S2211926417308676, doi: 10.1016/j.algal.2018.02.009.

Røkke GB, Hohmann-Marriott, MF Almaas E. An adjustable algal chloroplast plug-and-play model for genome-scale metabolic models. PLOS ONE. 2020 Feb; 15(2):e0229408. https://journals.plos.org/plosone/article?id=10.1371/journal.pone.0229408, doi: 10.1371/journal.pone.0229408, publisher: Public Library of Science.

Sayers EW, Bolton EE, Brister JR, Canese K, Chan J, Comeau D, Connor R, Funk K, Kelly C, Kim S, Madej T, Marchler-Bauer, A Lanczycki C, Lathrop S, Lu Z, Thibaud-Nissen, F Murphy T, Phan L, Skripchenko Y, Tse T, et al. Database resources of the National Center for Biotechnology Information. Nucleic Acids Research. 2021 Dec; 50(D1):D20–D26. https://www.ncbi.nlm.nih.gov/pmc/articles/PMC8728269/, doi: 10.1093/nar/gkab1112.

Scaife MA, Nguyen GTDT, Rico J, Lambert D, Helliwell KE, Smith AG. Establishing Chlamy-domonas reinhardtii as an industrial biotechnology host. The Plant Journal. 2015; 82(3):532–546. https://onlinelibrary.wiley.com/doi/abs/10.1111/tpj.12781, doi: 10.1111/tpj.12781, _eprint: https://onlinelibrary.wiley.com/doi/pdf/10.1111/tpj.12781.

Scranton MA, Ostrand JT, Fields FJ, Mayfield SP. Chlamydomonas as a model for biofuels and bio-products production. The Plant Journal. 2015; 82(3):523–531. https://onlinelibrary.wiley.com/doi/abs/10.1111/tpj.12780, doi: 10.1111/tpj.12780, _eprint: https://onlinelibrary.wiley.com/doi/pdf/10.1111/tpj.12780.

Yunus IS, Wichmann J, Wördenweber R, Lauersen KJ, Kruse O, Jones PR. Synthetic metabolic pathways for photobiological conversion of CO2 into hydrocarbon fuel. Metabolic Engineering. 2018 Sep; 49:201–211. https://www.sciencedirect.com/science/article/pii/S1096717618302210, doi: 10.1016/j.ymben.2018.08.008.

Yurkovich JT, Yang L, Palsson BO, Systems-level physiology of the human red blood cell is computed from metabolic and macromolecular mechanisms. bioRxiv; 2019. https://www.biorxiv.org/content/10.1101/797258v2, doi: 10.1101/797258, pages: 797258 Section: New Results.

Zhang Y, Pan Y, Ding W, Hu H, Liu J. Lipid production is more than doubled by manipulating a diacylglycerol acyltransferase in algae. GCB Bioenergy. 2021; 13(1):185–200.

Zheng HQ, Chiang-Hsieh, YF Chien CH, Hsu BKJ, Liu TL, Chen CNN, Chang WC. AlgaePath: comprehensive analysis of metabolic pathways using transcript abundance data from next-generation sequencing in green algae. BMC Genomics. 2014 Mar; 15(1):196. https://doi.org/10.1186/1471-2164-15-196, doi: 10.1186/1471-2164-15-196.

