## Supplementary figures and images for "Novel Context-Specific Genome-Scale Modelling Explores the Potential of *Chlamydomonas reinhardtii* for Synthetic Biology Applications"

### Supplemental Figures

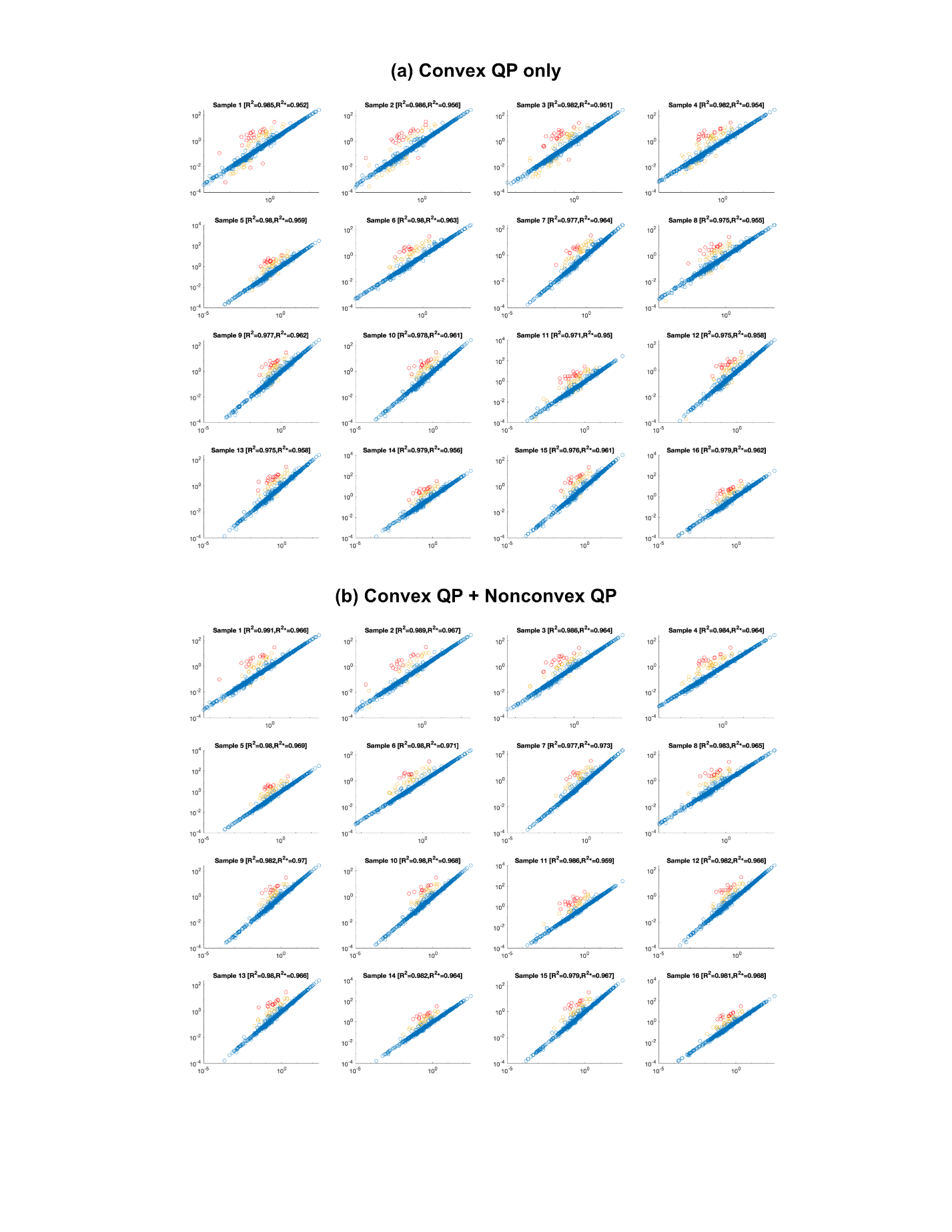

**Figure S1**

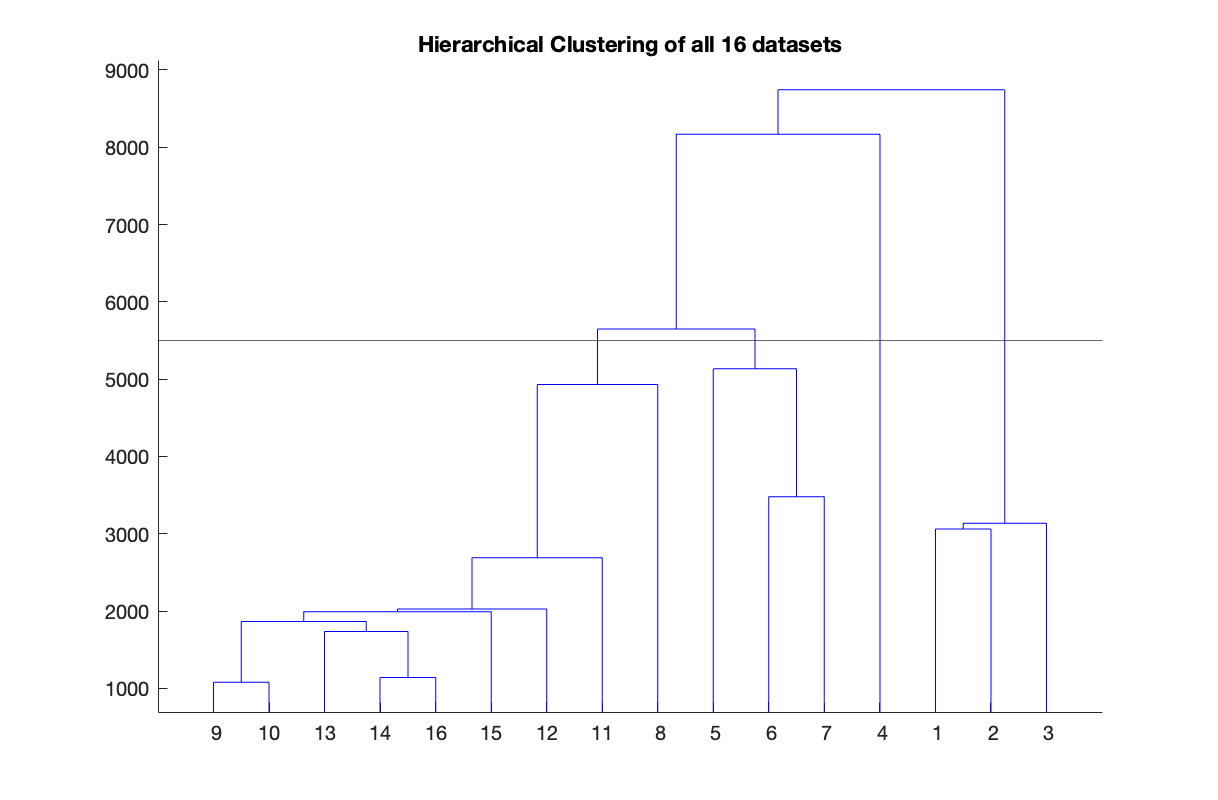

**Figure S2**

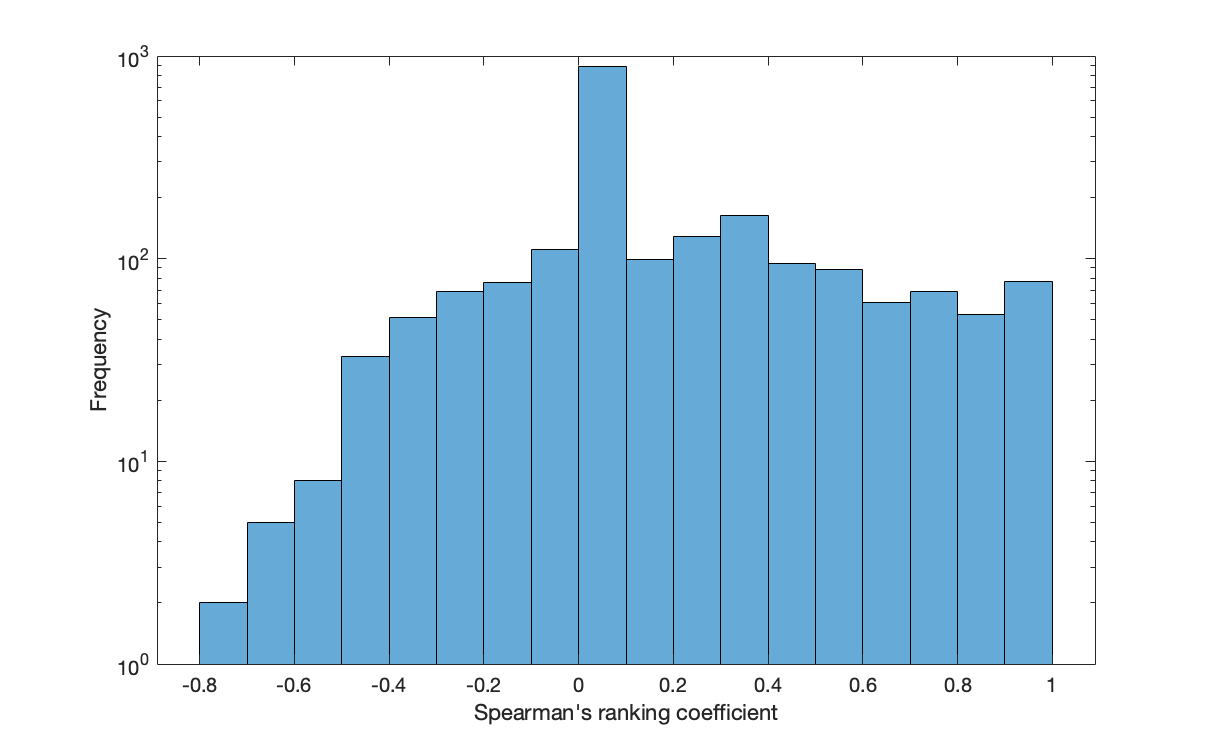
**Figure S3**
